# Identification of Ca-rich dense granules in human platelets using scanning transmission X-ray microscopy

**DOI:** 10.1101/622100

**Authors:** Tung X. Trinh, Sook Jin Kwon, Zayakhuu Gerelkhuu, Jang-Sik Choi, Jaewoo Song, Tae Hyun Yoon

## Abstract

Whole mount (WM) platelet preparations followed by transmission electron microscopy (TEM) observation is the standard method currently used to assess dense granule (DG) deficiency (DGD). However, due to electron density-based contrast mechanism in TEM, other granules such as α-granules might cause false DGs detection. Herein, scanning transmission X-ray microscopy (STXM), was used to identify DGs and minimize false DGs detection of human platelets. STXM image stacks of human platelets were collected at the calcium (Ca) L_2,3_ absorption edge and then converted to optical density maps. Ca distribution maps obtained by subtracting the optical density map at pre-edge region from those obtained at post-edge region were used for identification of DGs based on richness of Ca. Dense granules were successfully detected by using STXM method without false detection based on Ca maps for 4 human platelets. Spectral analysis of granules in human platelets confirmed that DGs contained richer Ca content than other granules. Image analysis of Ca maps provided quantitative parameters which would be useful for developing image-based DG diagnosis models. Therefore, we would like to propose STXM as a promising approach for better DG identification and DGD diagnosis, as a complementary tool to the current WM TEM approach.

## INTRODUCTION

Platelets are small and discoid-shaped anuclear blood cells, which are well known as essential components in hemostasis, the process of preventing blood loss through injured blood vessels. They contain several types of subcellular structures: α-granules, dense granules (DG), etc. Among these subcellular structures, deficiency of DGs, as found in previous research of Hermansky-Pudlak syndrome *(1)* and Chediak-Higashi syndrome *(2)*, would lead to bleeding diathesis. Evaluation of platelets by whole mount transmission electron microscopy (WM TEM) is an essential part in diagnostic workup of the dense granule deficiency (DGD) and is now considered the gold standard test for diagnosing DGDs *(3-10)*.

However, TEM has some drawbacks for the characterization of platelet DGs. Generally, WM TEM measurements were conducted without staining DGs, in which high contents of calcium (Ca) and phosphorus in platelet DGs are blocking electron beams and create contrast for DGs. However, TEM showed its best spatial resolution for the sample thickness less than ∼100nm and WM TEM observation of platelets with a variable thickness of 1-3μm often resulted in defocused and blurred images of subcellular granules. Additionally, because of the electron density-based contrast mechanism, other granules such as α-granules might cause false detection of DGs. Recently, scanning transmission X-ray microscopy (STXM) has shown its capabilities that may complement current limitations of WM TEM by providing better contrast mechanism and penetration depth with enough spatial resolution *(11-16)*. Particularly, the contrast mechanism of STXM, which is based on the differences in X-ray absorption at the pre- and post-edges, allows us to distinguish Ca-rich DGs, which may provide much clearer and more DG specific contrast than electron density-based contrast of TEM. However, STXM studies of platelets in the soft X-ray region have not been reported yet. In this study, synchrotron-based STXM at the soft x-ray region was adapted for the characterization of human platelet DGs and demonstrated its capability of identifying and counting DGs in human platelets and diagnosing DGD.

## RESULTS AND DISCUSSION

It has been known that DGs contain high contents of Ca and P *(4, 17-20)* making DGs electron-dense and allowing them to be visualized without special staining procedure. However, electron density-based contrast of the WM TEM is not element-specific and may have interference by the presence of other subcellular granules with high number density in platelets, such as alpha granules, and cause errors in counting DGs and diagnosing DGDs.

The high content of Ca in DG was also chosen in this STXM study to distinguish DGs from other subcellular granules. However, in contrast with the electron density-based contrast mechanism of the TEM, the STXM approach uses a more distinct contrast mechanism based on element- or chemical species-specific XANES (X-ray absorption near edge structure) features, which allows us much clearer identification of platelet DGs. As shown in Figure 1, STXM image stack of a whole mounted platelet was collected and converted to OD maps. Then, Ca distribution maps was obtained by subtracting the OD map at the pre-edge region from the OD map at the post-edge region. In the STXM images collected at the energy of 345.6eV and 349.3eV (Figure 1A and 1C), many subcellular granules were observed with good contrast. When these images were converted to OD maps, one granule became even brighter in the OD image collected at 349.3eV (Figure 1D), while the other granules remained with similar contrast (Figure 1B and 1D). In the Ca distribution map shown in Figure 1E, only one granule was observed as a bright spot, while the others were mostly dimmed. This bright spot in Figure 1E is the Ca-rich sub-cellular granule of platelets, which can be clearly identified as DGs.

**Figure 1.**
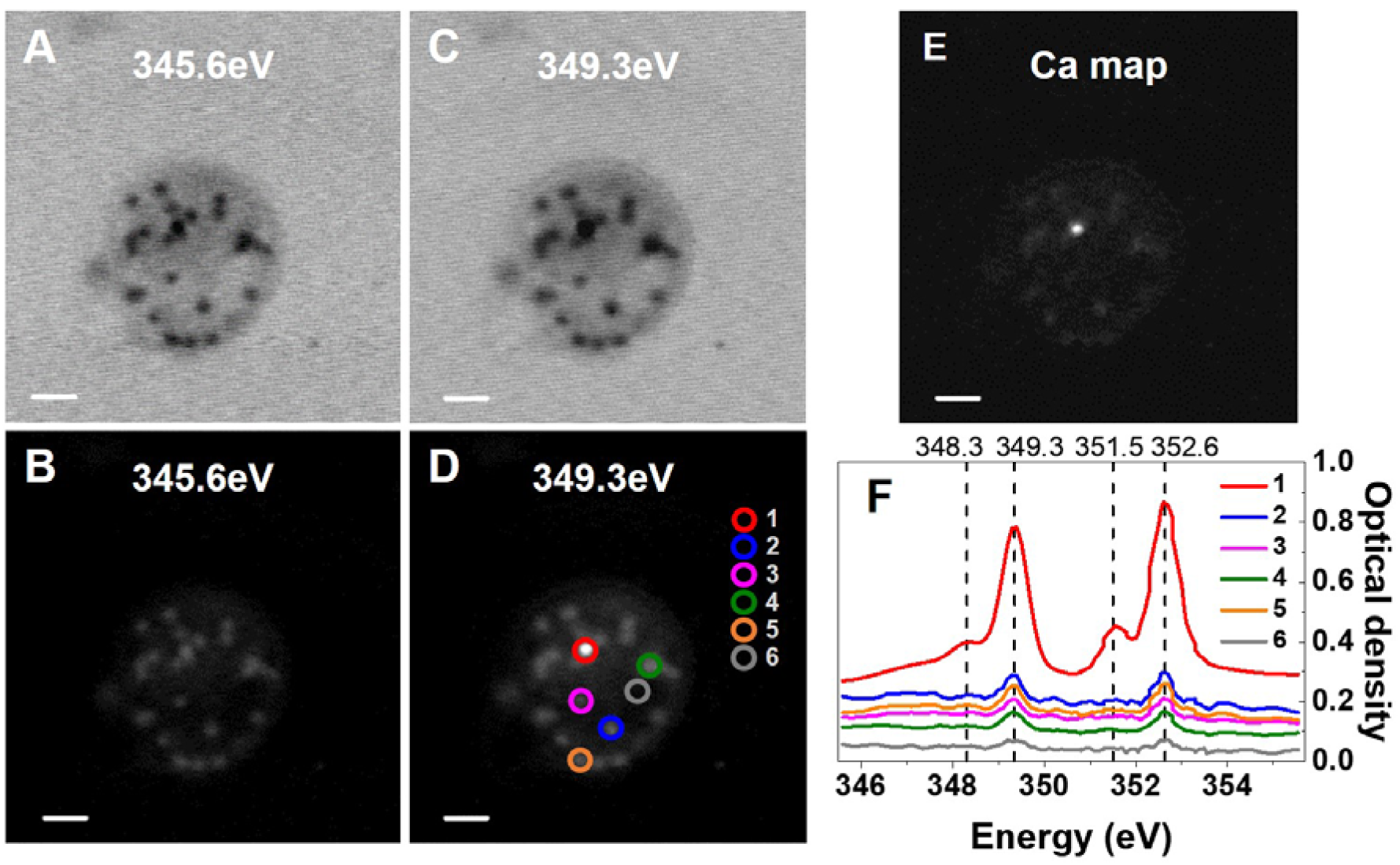
STXM transmission images at 345.6eV (A) and 349.3eV (C), STXM OD images at 345.6eV (B) and 349.3eV (D) of platelet 1; Calcium map (E) of platelet 1 obtained by subtracting B from D; XANES spectra (F) of six regions inside platelet 1 (marked in D). Scale bars are 1µm.

Additionally, six different regions of the platelet (regions 1-6 of Figure 1D) were chosen and their Ca L_2,3_ edge XANES spectra were displayed in Figure 1F. The XANES spectrum of the brightest spot in the Ca map (region 1 of Figure 1D) displayed typical Ca L_2,3_ edge XANES spectrum, similar with those previously reported in the literature *(13, 21-23)*, with two strong (349.3eV and 352.6eV) and two weak (348.3eV and 351.5eV) absorption peaks. The XANES spectra of the other four Ca-poor granules (regions 2-5 of Figure 1D) also displayed similar spectral features with much reduced intensities, indicating that these granules contain much less amount of Ca. Moreover, the smaller peaks at 348.3eV and 351.5eV seem almost disappeared, which might be due to the presence of different Ca species (e.g., Ca-carbonates, Ca-phosphates, biogenic Ca etc.) in these Ca-rich and Ca-poor granules. A small absorption peak observed for the cytoplasm region of the platelet suggest that only a negligible amount of Ca was present in the cytoplasm of the platelet (region 6 of Figure 1D). With these Ca L_2,3_ edge XANES spectra and the Ca distribution map, we can confirm that the brightest spot in the Ca map (Figure 1E) was a DG, while the other particles, which were observed as bright spots in the OD map at 349.3eV but found as dimmed in the Ca maps, were probably other types of subcellular granules.

Further collection of STXM images were performed for three other platelets (platelet #2-#4) and their original and processed STXM images were presented in Figure 2. In the Ca distribution maps of these platelets, DGs are clearly distinguished as bright spots within the cytoplasm of platelets (Figure 2, A5– D5). Further image analysis of Ca maps of these platelets allows us to extract additional parameters for dense granules such as area, circularity, diameters, and Ca content for each granule (Table 1). These parameters should be valuable for further developing image-based DG diagnosis models as well as image interpretation criteria. Recently, Chen et al. *(10)* proposed the DG interpretation criteria for WM TEM images, in which typical microscopic features of true DGs were described as uniformly dense/dark textures, perfectly round shape with sharp contour and larger than 50nm in diameter. As a complementary approach of WM TEM, the STXM based Ca distribution map may provide additional DG interpretation criteria that may refine and improve accuracy of DGD diagnosis. Although follow-up studies with larger number of platelets are required for statistically meaningful interpretation of these STXM observations, we think that our study have shown the potential capabilities of STXM to identify and count DGs as a complementary method of typical WM TEM approach. Furthermore, the STXM also have the potentials for semi-quantitative Ca contents estimation and chemical species-specific maps, which may further improve accuracy of DGD diagnosis.

**Table 1.** Image analysis results for the dense granules in STXM-based Ca maps for platelets #1 - #4.

| Platelet | Dense granules identified | Area (µm<sup>2</sup>) | Circularity | Feret diameter (µm) | Diameter based on circle shape (µm) | Average intensity in Ca Map | Maximum intensity in Ca Map |
| --- | --- | --- | --- | --- | --- | --- | --- |
| #1 | 1 | 0.065 | 0.917 | 0.329 | 0.288 | 0.590 ± 0.294 | 1.137 |
| #2 | 1 | 0.024 | 0.904 | 0.216 | 0.175 | 0.176 ± 0.099 | 0.438 |
|  | 2 | 0.026 | 0.897 | 0.247 | 0.182 | 0.173 ± 0.095 | 0.390 |
|  | 3 | 0.018 | 0.897 | 0.204 | 0.151 | 0.224 ± 0.107 | 0.336 |
|  | 4 | 0.021 | 1.000 | 0.204 | 0.164 | 0.232 ± 0.127 | 0.487 |
| #3 | 1 | 0.051 | 0.572 | 0.351 | 0.255 | 0.316 ± 0.128 | 0.560 |
|  | 2 | 0.026 | 0.624 | 0.224 | 0.182 | 0.305 ± 0.083 | 0.487 |
| #4 | 1 | 0.024 | 0.944 | 0.199 | 0.175 | 0.326 ± 0.117 | 0.506 |

**Figure 2.**
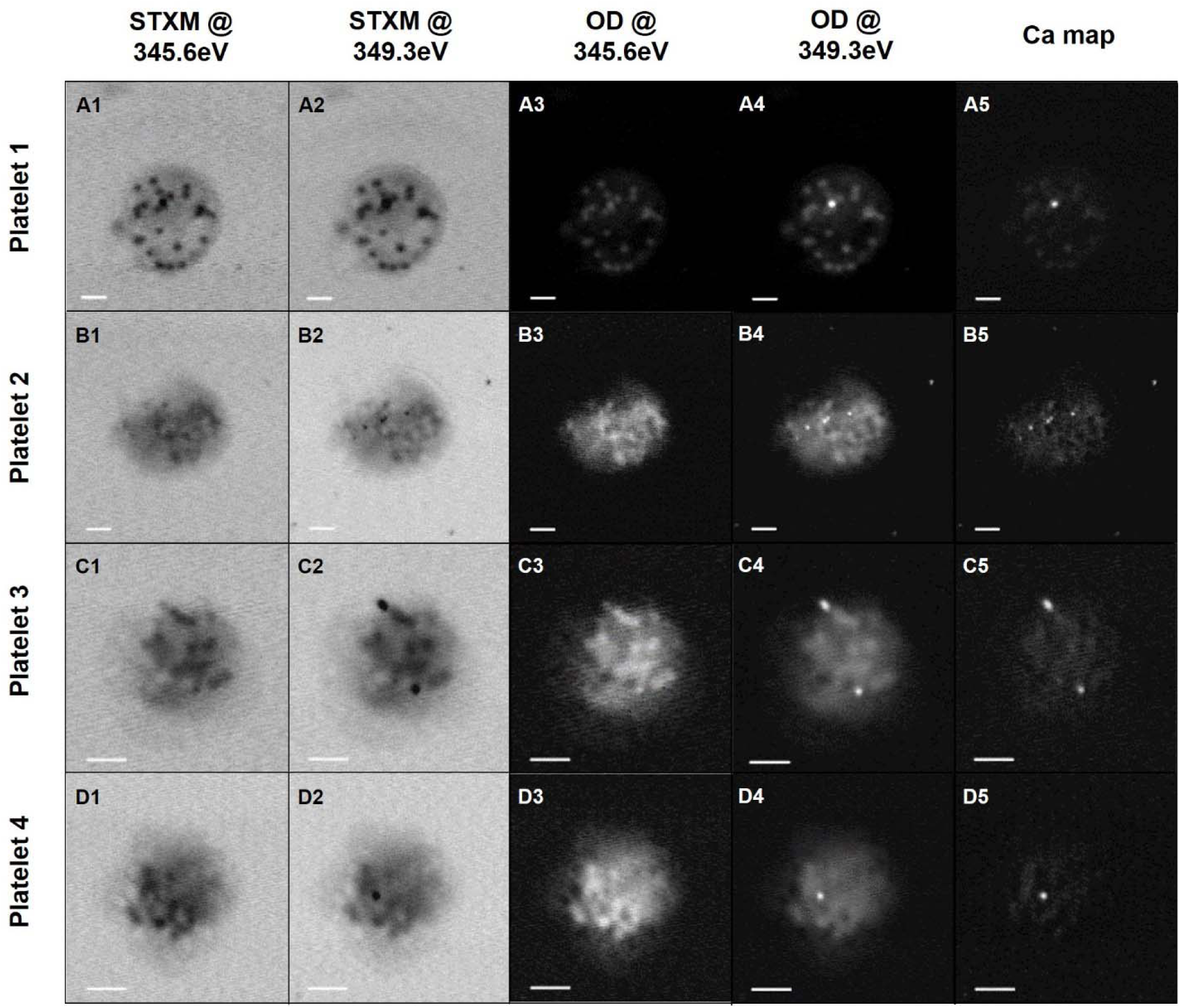
STXM transmission images at 345.6eV (A1 – D1) and 349.3eV (A2 – D2); STXM OD images at 345.6eV (A3 – D3) and 349.3eV (A4 – D4) and Calcium maps (A5 – D5) of four platelets 1 – 4. Scale bars are 1µm.

In conclusion, X-ray spectromicroscopic technique (i.e., STXM) was adapted to identify and count DGs of human platelets and demonstrated its promising capability in identifying and counting DGs in human platelets. Element-specific contrast mechanism based on the differences in X-ray absorption edge allow us to distinguish Ca-rich DGs more clearly from the other subcellular components. We propose STXM as a promising approach for DGD diagnosis with an improved identification and counting capability, which complement the current WM TEM method.

## METHODS

### Platelet Samples Preparation

Whole blood was obtained from normal healthy persons after informed consent and the study was approved by the Institutional Review Board of Yonsei Severance Hospital, Seoul. Fresh Platelet-rich plasma (PRP) was used for STXM measurement. Whole blood was drawn into tubes containing acid citrate dextrose (one-sixth the volume of blood drawn). PRP was obtained by centrifugation at 210 x g for 15 minutes at 37°C. Before preparing STXM samples, fresh PRP was washed using deionized water 5 times with the following protocol of adding 800µL of deionized water then centrifuging and removing 800 µL of the upper solution. A single drop of the remaining platelet solution (approximately 2 µL) was transferred to the Formvar-coated copper grid following by air-drying overnight. The excess solution of fresh platelets was removed by using filter paper. The copper grids containing dried platelets were then used for STXM measurements.

### STXM measurement

STXM images were acquired at the 10A beamline of the Pohang light source (PLS, Pohang Accelerator Laboratory, Republic of Korea). The storage ring current of 200 mA at the PLS was operated in top-up mode. The synchrotron-based monochromatic soft X-ray was focused to ∼25 nm using the Fresnel zone plate (outer most zone width of 25 nm). The first order of a diffractive focused X-ray was selected using the order-sorting aperture (OSA, pinhole diameter of 100 µm). The sample was mounted on the interferometrically controlled piezo stage and raster scanned. Intensity of the transmitted X-ray was measured by using a scintillator photomultiplier tube (PMT). STXM images of whole mounted platelets were collected for the area of 10µm × 10µm with 300×300 pixels. To obtain Ca L_2,3_ edge XANES spectra, stack images of platelets were collected at the energy range of 345.6 – 355.6 eV. The energy position of the Ca L_3_-edge and L_2_-edge main peaks were calibrated to 349.3 eV and 352.6 eV, respectively, according to Rieger et al *(24)*. The aXis2000 software *(25)* (version 01-Oct-2018) was used to calculate the optical density (OD) and Ca distribution maps.

## Acknowledgment

This work was supported by the Bio & Medical Technology Development Program of the National Research Foundation (NRF) funded by the Ministry of Science & ICT (No. 2017M3A9G8084539). This research was also partially supported by the National R&D Program through the National Research Foundation of Korea (NRF) funded by the Ministry of Science, ICT and Future Planning (2017K1A3A7A09016394). We also would like to thank Dr. Namdong Kim, Dr. Hyunjun Shin and Ms. Mikang Kim of Pohang Light Source for their assistance at the STXM beamline.

## Author Contributions

T.H.Y. designed and supervised research. T.X.T. and S.J.K. performed the STXM experiment, Z.G. and J.S. prepared platelet samples, T.X.T. and J.S.C. performed the STXM image analysis and T.X.T. and T.H.Y. wrote manuscript.

## Additional Information

